# Integrated Bayesian analysis of rare exonic variants to identify risk genes for schizophrenia and neurodevelopmental disorders

**DOI:** 10.1101/135293

**Authors:** Hoang T Nguyen, Julien Bryois, April Kim, Amanda Dobbyn, Laura M Huckins, Ana B Munoz-Manchado, Douglas M Ruderfer, Giulio Genovese, Menachem Fromer, Xinyi Xu, Dalila Pinto, Sten Linnarsson, Matthijs Verhage, August B Smit, Jens Hjerling-Leffler, Joseph Buxbaum, Christina Hultman, Pamela Sklar, Shaun M Purcell, Kasper Lage, Xin He, Patrick F Sullivan, Eli A Stahl

**Author notes:** Co-correspondence author, Division of Psychiatric Genomics, Department of Genetics and Genomic Sciences, Institute for Genomics and Multiscale Biology, Icahn School of Medicine at Mount Sinai, New York, New York, 10029, USA.

## Abstract

**Background:** Integrating rare variation from trio family and case/control studies has successfully implicated specific genes contributing to risk of neurodevelopmental disorders (NDDs) including autism spectrum disorders (ASD), intellectual disability (ID), developmental disorders (DD), and epilepsy (EPI). For schizophrenia (SCZ), however, while sets of genes have been implicated through study of rare variation, only two risk genes have been identified.

**Methods:** We used hierarchical Bayesian modeling of rare variant genetic architecture to estimate mean effect sizes and risk-gene proportions, analyzing the largest available collection of whole exome sequence (WES) data for schizophrenia (1,077 trios, 6,699 cases and 13,028 controls), and data for four NDDs (ASD, ID, DD, and EPI; total 10,792 trios, and 4,058 cases and controls).

**Results:** For SCZ, we estimate 1,551 risk genes, more risk genes and weaker effects than for NDDs. We provide power analyses to predict the number of risk gene discoveries as more data become available, demonstrating greater value of case-control over trio samples. We confirm and augment prior risk gene and gene set enrichment results for SCZ and NDDs. In particular, we detected 98 new DD risk genes at FDR *<* 0.05. Correlations of risk-gene posterior probabilities are high across four NDDs (*ρ >* 0.55), but low between SCZ and the NDDs (*ρ <* 0.3). In depth analysis of 288 NDD genes shows highly significant protein-protein interaction (PPI) network connectivity, and functionally distinct PPI subnetworks based on pathway enrichments, single-cell RNA-seq (scRNAseq) cell types and multi-region developmental brain RNA-seq.

**Conclusions:** We have extended a pipeline used in ASD studies and applied it to infer rare genetic parameters for SCZ and four NDDs. We find many new DD risk genes, supported by gene set enrichment and PPI network connectivity analyses. We find greater similarity among NDDs than between NDDs and SCZ. NDD gene subnetworks are implicated in postnatally expressed presynaptic and postsynaptic genes, and for transcriptional and post-transcriptional gene regulation in prenatal neural progenitor and stem cells.

## Background

Integrating rare variation from family and case-control studies has successfully implicated specific genes contributing to risk of neurodevelopmental disorders (NDDs) including autism spectrum disorders (ASD), intellectual disability (ID), developmental disorders (DD) and epilepsy (EPI). These early-onset disorders typically manifest as infant or childhood developmental delay or regression, and can be comorbid even within individuals [1] at the symptom and syndrome levels. ASD typically includes deficits in social function and often includes cognitive deficits; ID is denfined by severe cognitive deficits; DD is characterized by physical or neurological developmental delays delays frequently including ID; and EPI is defined by recurrent seizures and often occurs in probands of the other NDDs [2, 3, 4]. Cognitive dysfunction is a common thread among these disorders and many of the risk genes identified for them point to brain neuronal development as well as synaptic function.

For schizophrenia (SCZ), however, while sets of genes have been implicated through study of rare variation (including NDD risk genes) [5, 6, 7], only two risk genes containing rare exonic variants of strong effect have been identified [6, 8, 9]. SCZ is an etiologically complex psychiatric disorder characterized by hallucinations, delusions, and cognitive symptoms; heritability is estimated to be 60-80% [10, 11]; and the genetic architecture of SCZ is highly polygenic with contributions of common variation and rare inherited and *de novo* structural and exon variants [12, 13, 5, 6, 14, 8, 7, 15]. With the advent of affordable high-quality next-generation sequencing, the genetics of SCZ and other diseases can be increasingly better characterized, especially for rare variants. Rare variants in case-control and trio samples have been leveraged to identify SCZ genes and gene sets. However, SCZ rare variant genetic architecture remains poorly understood. Such analyses could help gain further insights into this disease, for example by using the estimated number of risk genes to calibrate gene discovery false discovery rates, or by using the distribution of effect sizes to improve power estimates and rare variant association study design. A better understanding of our certainty in sets of risk genes for SCZ will provide a better picture of biological pathways relevant for the disease.

We developed an improved hierarchical Bayesian modeling framework [16], extended Transmission And De novo Association, `extTADA`, to analyze whole exome sequence (WES) data in SCZ and four NDDs (ASD, ID, DD, EPI), which have substantial clinical and etiological overlap. All are brain diseases with prominent impacts on cognitive function. Multiple recent studies supporting genetic overlap among these disorders have included common variant genetic correlations [17, 18] and shared molecular pathways [19, 20], and shared genes with *de novo* mutations [6, 21]. Using the largest sample assembled to date for a unified analysis of these disorders, we find greater overlap among the NDDs than with SCZ, despite the emphasis on overlap in the SCZ rare variant literature [6, 19, 7]. We used `extTADA` statistical support to compile a comprehensive list of 288 NDD genes. Network analyses of these genes begin to pinpoint and intersect functional processes implicated in disease, brain cell types and developmental timepoints of expression.

## Data and methods

### Data

Figure S1 shows the workflow of all data used in this study.

*Variant data of SCZ, ID, DD, EPI and ASD*

High-quality variants were obtained from published analyses (Table S1). For SCZ case-control data, we performed exome-wide association analyses to test for stratification and to identify non-heterogeneous samples for `extTADA` analysis (see Supplementary Methods).

Variants were annotated using Plink/Seq (using RefSeq gene transcripts, UCSC Genome Browser [22]) as described in Fromer at al., 2014 [6]. SnpSift version 4.2 [23] was used to further annotate these variants using dbnsfp31a [24]. Variants were grouped into different categories as follows: loss of function (LoF, nonsense, essential splice, and frameshift variants); missense damaging (MiD, defined as missense by Plink/Seq and damaging by all of seven methods [7]-SIFT, *P olyphen*2 *HDIV*, *P olyphen*2 *HV AR*, LRT, PROVEAN, MutationTaster and MutationAssessor); synonymous mutations in regulatory regions as described in [25]. To annotate synonymous variants within DNase I hypersensitive sites (DHS) as [25], the file *wgEncodeOpenChromDnaseCerebrumfrontalocPk.narrowPeak.gz* was downloaded from [26] on April 20, 2016. Based on previous results with SCZ exomes [5, 7], only case-control singleton variants were used in this study (i.e., observed once). The data from Exome Aggregation Consortium (ExAC) [27] were used to annotate variants inside ExAC (InExAC or not private) and not inside ExAC (NoExAC or private). On April 20, 2016, the file *ExAC.r0.3.nonpsych.sites.vcf.gz* was downloaded from [28] and BEDTools was used to obtain variants inside (InExAC) or outside this file (NoExAC).

#### Mutation rates

We used the methodology which was based on trinucleotide context[29, 30], and incorporating depth of coverage [6], to obtain mutation rates for each variant annotation category. We assigned one tenth the minimum non-zero mutation rate to genes with calculated mutation rates equal to zero.

#### Gene sets

Multiple resources were used to obtain gene sets for our study. First, we used known and candidate gene sets with prior evidence for involvement in SCZ and ASD. Second, to identify possible novel significant gene sets, we collected genes sets from available data bases (see below).

#### Known/candidate gene sets

These gene sets and their abbreviations are presented in Table S2. They included: gene sets enriched for ultra rare variants in SCZ which were described in detailed in [7] consisting of missense constrained genes (constrained) from [29], loss-of-function intolerant genes (pLI90) from [27], *RBFOX2* and *RBFOX1/3* target genes (rbfox2, rbfox13) from [31], Fragile X mental retardation protein target genes (fmrp) from [32], *CELF4* target genes (celf4) from [33], synaptic genes (synaptome) from [34], microRNA-137 (mir137) from [35], PSD-95 complex genes (psd95) from [36], ARC and NMDA receptor complexes (arc, nmdar) genes from [37], *de novo* copy number variants in SCZ, ASD, bipolar as presented in Supplementary Table 5 of [7]; allelic-biased expression genes in neurons from Table S3 of [38]; promoter targets of *CHD8* from [39]; known ID gene set was from the Sup Table 4 of [40] and the 10 novel genes reported by [40]; gene sets from MiD and LoF *de novo* mutations of ASD, EPI, DD, ID; the essential gene set from the supplementary data set 2 of [41]; lists of human accelerated regions (HARs) and primate accelerated regions (PARs) [42] were downloaded from [43] on May 11, 2016 (the coordinates of these regions were converted to hg19 using Liftover tool [44]. We used a similar approach as [45] to obtain genes nearby HARs. Genes in regions flanking 100 kb of the HARs/PARs were extracted to use in this study, geneInHARs, geneInPARs); list of known epilepsy genes was obtained from Supplementary Table 3 of [46]; list of common-variant genes was obtained from Extended Table 9 of [15]; list of gene sets from 24 modules from Supplementary Table 2 of [47]; 134 gene sets from mouse mutants with central nervous system (CNS) phenotypes were obtained from [15, 48].

In the gene-set tests for a given disease, we removed the list of known genes and the list of *de novo* mutation genes for that disease. As a result, we tested 185 candidate gene sets for ASD, DD and SCZ; and 184 candidate gene sets for EPI and ID.

#### Other gene sets

We also used multiple data sets to identify novel gene sets overlapping with the current gene sets. Gene sets from the Gene Ontology data base [49], KEGG, REACTOME and C3 motif gene sets gene sets collected by the Molecular Signatures Database (MSigDB) [50]. To increase the power of this process, we only used gene sets with between 100 to 4995 genes. In total, there were 1717 gene sets. These gene sets and the above gene sets were used in this data-driven approach.

#### Transcriptomic data

Spatio-temporal transcriptomic data were obtained from BRAINSPAN [51]. The data were divided into eight developmental time points (four pre-natal and four postnatal) [52]. Single-cell RNAseq data were obtained from [53].

### Methods

#### The extTADA pipeline

We extended the hierarchical Bayesian modeling framework of [16] to develop extended Transmission And De novo Association, `extTADA` (Figure S2, Table S3) for Bayesian analysis via Markov Chain Monte Carlo (MCMC). Primary extensions to the TADA model [16, 30] for parameter inference include fully joint inference, approximating the CC data probability so that population allele frequency parameters were eliminated, and constraining effect size distribution to prevent an implied proportion of protective variants.

Additional details are described in Supplementary Methods, and at *https://github.com/hoangtn/extTADA*. Briefly, for a given gene, all variants of a given category (e.g. either *de novo* or singleton case-control loss-of-function) were collapsed and considered as a single count. Let *γ* be the mean relative risk (RR) of the variants, which is assumed to follow a distribution across risk genes. The data likelihood was considered a mixture of non-risk and risk gene hypotheses, *H*_0_ : *γ* = 1 and *H*_1_ : *γ ≠* 1,

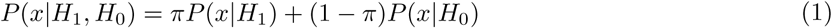

where *H*_0_ and *H*_1_ represent *γ* and all other parameters under the model, and the mixture proportion *π* is interpreted as the proportion of risk genes genome wide.

The data *x* are DN and CC counts (*x*_*dn*_, *x*_*ca*_, *x*_*cn*_), and the `extTADA` likelihood was the product of data probabilities over any numbers of population samples and variant categories. Hyper parameters for different categories in Table S3 were jointly estimated based on the mixture model,

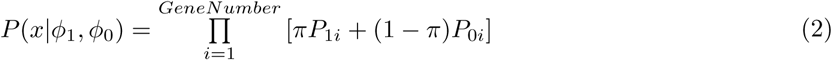

where *P*_1*i*_ and *P*_0*i*_ at the *i*^*th*^ gene were calculated across population samples and categories as follows:

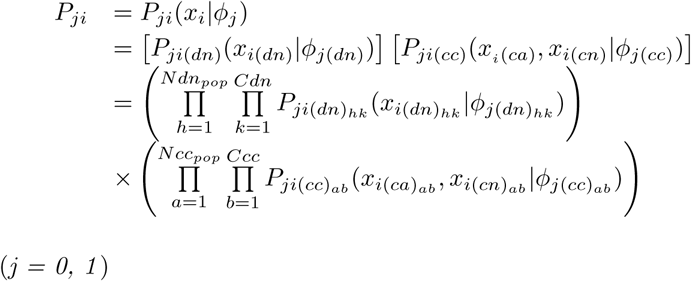

(*j = 0, 1*)

To simplify the estimation process in Equation 2, we approximated the original `TADA` model for CC data *P* (*x*_*ca*_, *x*_*cn*_*|H*_*j*_) using a new model in which case counts were conditioned on total counts: *P* (*x*_*ca*_*|x*_*ca*_ + *x*_*cn*_, *H*_*j*_) (Figure S2 and Supplementary Methods).

`extTADA` used Markov Chain Monte Carlo (MCMC) for Bayesian analysis. We extracted posterior density samples from at least two MCMC chains. Posterior modes were reported as parameter estimates for all analyses, with 95% credible intervals (CIs).

Then, gene-level Bayes Factors (*BF*_*gene*_) can be calculated for each category to compare hypotheses *H*_1_ and *H*_0_ (*BF* = *P* (*x|H*_1_)*/P* (*x|H*_0_)). *BF*_*gene*_ could be calculated as the product of BFs across multiple variant categories. Data could be from heterogeneous population samples; therefore, we extended `TADA`’s *BF*_*gene*_ as the product of BFs of all variant categories including population samples as in Equation 3,

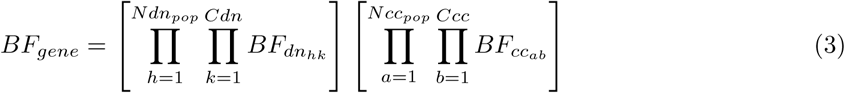

in which *N dn*_*pop*_, *N cc*_*pop*_ are the numbers of DN and CC population samples, and *C*_*dn*_, *C*_*cc*_ are the number of annotation categories in DN and CC data. We changed the order of integrals to avoid numerical integration over *P* (*q*) because the true range of this parameter is not known (Supp Methods). We inferred significant genes by converting BFs to false discovery rates (FDRs) using the approach of [54] as described in [30]. Posterior probability (PP) for each gene was calculated as *P P* = *π * BF/*(1 *- π* + *π * BF*) [55].

#### Testing the pipeline on simulated data

To test `extTADA`, we used the simulation method described in the `TADA` paper [16]. We simulated one case-control (CC) variant class, two CC classes, or one CC and one *de novo* (DN) class. For CC data, the original case-control model in TADA [16] was used to simulate case-control data and then case-control parameters were estimated using the approximate model. The frequency of SCZ case-control LoF variants was used to calculate the prior distribution of *q ∼ Gamma*(*ρ, ν*) as described in Table S3.

Different sample sizes were used. For CC data, to see the performance of the approximate model, we used four sample sizes: 1092 cases plus 1193 controls, 3157 cases plus 4672 controls, 10000 cases plus 10000 controls, 20000 cases plus 20000 controls. The first two sample sizes were exactly the same as the two sample sizes from Sweden data in current study. The last two sample sizes were used to see whether the model would perform better if sample sizes increased. For DN and CC data, we used exactly the sample sizes of the largest groups in our current data sets: family numbers = 1077, case numbers = 3157 and control numbers = 4672.

To assess the performance of model parameter estimation, we calculated Spearman correlation coefficients [56] between estimated and simulated parameter values. For each combination of simulated parameters, we re-ran 100 times and obtained the medians of estimated values to use for inferences. We also used different priors of hyper parameters (e.g., 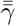, 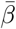 in Table S3) in the simulation process and chose the most reliable priors corresponding with ranges of 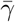. Because 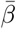 mainly controlled the dispersion of hyper parameters, 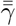 was set equal to 1, and only 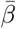 was tested.

To assess the performance of `extTADA` risk gene identification, we compared expected and observed FDRs (oFDRs). We defined oFDR as the proportion of FDR significant genes that were true risk genes (determined for data simulation). We simulated DN and CC data for a range of sample sizes, using parameter values randomly sampled from the posterior density of our primary SCZ analysis.

We also conducted power analyses of larger sample SCZ studies using parameters sampled from the posterior density of our primary SCZ analysis. For power analyses, we assumed sample sizes ranging from 500 to 20,000 trio families and equal numbers of cases and controls ranging from 1000 to 50,000 of each, and calculated the number of risk genes at FDR *≤* 0.05.

We also tested the situation in which no signal of both *de novo* mutations and rare case-control variants was present. We simulated one DN category and one CC category with *π* = 0, 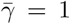. To see the influence of prior information of 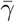 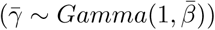 on these results, we used different values of 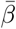.

#### Applying extTADA to real data

*Estimating genetic parameters* For SCZ, DN mutations and CC variants from non-heterogeneous population samples were analyzed. Three DN mutation categories (MiD, LoF, and silentFCPk mutations) and one CC variant category (MiD and LoF variants, pooled) were used in Equation 2 to obtain genetic parameters for SCZ. For ASD, two DN (MiD, LoF) and one CC (MiD and LoF pooled) variant categories were analyzed. For the three other disorders, only *de novo* data (MiD, LoF categories) were analyzed because there were not rare case-control data available.

#### Secondary analyses

We compared our results with those generated using mutation rates adjusted for the ratio of observed to expected synonymous mutations. We divided the observed counts by expected counts (= 2 x family numbers * total mutation rates), and then used this ratio to adjust for all variant category mutation rates.

We conducted further analyses of the SCZ data. Each variant category (LoF, MiD, silentFCPk DN mutations, LoF+MiD CC variants) was analyzed individually to assess their contributions to the primary results. We conducted secondary analyses including CC variants present in ExAC, and with equal mean RR parameters (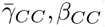) across CC population samples.

#### Gene set enrichment in extTADA results

Based on the `extTADA` results, we tested the enrichment of gene sets by using gene PPs as follows. At each gene, we obtained PP from `extTADA`. For each tested gene set, we calculated the mean of PPs (*m*_0_). After that, we randomly chose gene sets matched for mutation rates and recalculated mean PP *n* times (n = 10 million in this study) (generating the vector *m*). The empirical p value for the gene set was calculated as: 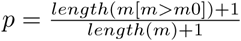. To correct for multiple tests, the p values were FDR adjusted using the method of [57]. To match mutation rates, for each gene we chose random genes from the 1000 genes with closest mutation rates.

To test the results of the mean-PP based method above, we also compared the method with a permutation based method. For each condition, we chose the top 500 genes with smallest FDR values from `extTADA` results. For each gene set, we calculated the number of overlapping genes between the 500 genes and the gene set (*m*_0_). After that, we randomly chose gene sets having the same length as the tested gene set, and recorded the intersecting gene number with the top 500 genes. This process was carried out n times to have a vector *m* (n = 10,000,000). Matching genes by mutation rate and empirical p value calculation were as described above.

#### pLI/RVIS data in novel significant gene sets

Residual Variation Intolerance Score (RVIS) information (*RVIS Unpublished ExACv2 March2017.txt*) was downloaded from [58] and probability of loss-of-function intolerant (pLI) information was downloaded from [59], on June 20 2017. To calculate p, *µ, σ, z* values for a gene set, we used the same approach as [40] with 10,000 permutations.

#### Single cell enrichment analysis

We obtained gene expression from 9970 single cells that were previously clustered into 24 different cell types [53]. We used the `scran` R package [60, 61] using the 50% of the genes with mean expression higher than the median to compute normalization factor for each single cell. The normalization factors were computed after clustering cells using the `scran` *quickcluster()* function to account for cell type heterogeneity. We then performed 24 differential expression analysis using BPSC [62] testing each cell type against the 23 other cell types using the normalization factors as covariates. For each differential expression analysis, the t-statistics were then standard normalized. Finally, for each cell type, we tested if the standard normalized t-statistic for genes in the gene sets were significantly higher than genes not in the gene set.

#### Network and transcriptome analyses

We used GeNets [63] to test protein interactions from gene sets. Connectivity P values were obtained by permuting 75,182 matched random networks, and communities (subnetworks showing greater connectivity within than between) were defined by hierarchical agglomeration [64]. Spatiotemporal transcriptome data were clustered using a hierarchical method inside the *heatmap.2* of the package gplots [65]. We used a height = 9 (in the function *cutree*) to divide data into 8 groups from clustering results. Default options were used for this clustering process. Fisher’s exact test [66] was used to obtain p values between spatiotemporal transcriptome clusters and GeNets based communities.

## Results

### The extTADA pipeline for rare variant genetic architecture inference

We first present a pipeline for integrative analysis of trio-based *de novo* (DN) variants and case-control (CC) rare variants, in order to infer rare variant genetic architecture parameters and to identify disease risk genes. We extended the hierarchical Bayesian modeling framework of He *et al.* [16] to develop extended Transmission And De novo Association, `extTADA` (Figure S2, Table S3) for Bayesian analysis via Markov Chain Monte Carlo (MCMC). Primary extensions to the TADA model [16, 30] for parameter inference include fully joint inference, eliminating population allele frequency parameters in parameter inference, and constraining effect size distributions to prevent an implied proportion of protective variants.

Very briefly, variants are collapsed and considered as a single integer count per gene with relative risk *γ*. The likelihood under a mixture model of risk gene and non-risk gene hypotheses, *H*_0_: *γ* = 1 and *H*_1_: *γ ≠* 1, is 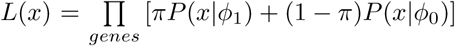, where *ϕ*_0,1_ represent all model parameters (Table S3). Multiple variant categories and population samples are incorporated as in Eq. 2. MCMC posterior sample modes are reported as parameter estimates with 95% credible intervals (CIs). The key parameters are 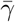, the mean relative risk across risk genes, and the mixture proportion *π*, which is interpreted as the proportion of risk genes genome wide and allows estimation of the number of disease risk genes.

Bayes factors, *BF* = *P* (*x|ϕ*_1_)*/P* (*x|ϕ*_0_), quantify support for each gene being a risk gene, and are converted to posterior probabilities [55] and frequentist false discovery rates (FDRs) [54, 30] for ease of interpretation.

#### Evaluating extTADA on simulated data

We analyzed simulated DN and CC data with one variant category each and CC data with two variant categories, to examine inference on a single variant class as well as to assess the conditional probability approximation for CC data (Figures S4, S5, S6 and S7, Supplementary Results). We tested sample sizes ranging from that of the available data, 1,077 trios and 3,157 cases (equal controls), and larger sample sizes of up to 20,000 cases (see Supplementary Results).

We observed little bias in parameter estimation (Table S4 and S5). With very large relative risks of the inherited variants, we observed slight under and over estimation of the risk gene proportion and RR, respectively; we note that these conditions appear outside the range of our SCZ analyses. Some bias can be expected in Bayesian analysis and does not have a large effect on the risk gene identification under this model [16]. We assessed this directly by calculating observed FDR (oFDR, i.e. the proportion of genes meeting a given FDR significance threshold that are true simulated risk genes). `extTADA` risk gene identification results were well-calibrated (Figure 1) over wide parameter ranges. For small *π* (e.g., *π* = 0.02), oFDRs were higher than FDRs when *de novo* mean RRs were small (*∼* 5). We also observed oFDR were equal to zero for some cases with small FDR, when very small numbers of FDR-significant genes were all true risk genes. We also ran `extTADA` on null data, *π* = 0 and 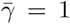 for both DN and CC data (Table S6); here, MCMC chains tended not to converge, *π* estimates trended to very small values, and Bayes factors and FDRs identified almost no FDR-significant genes as expected (Table S6).

**Figure 1.**
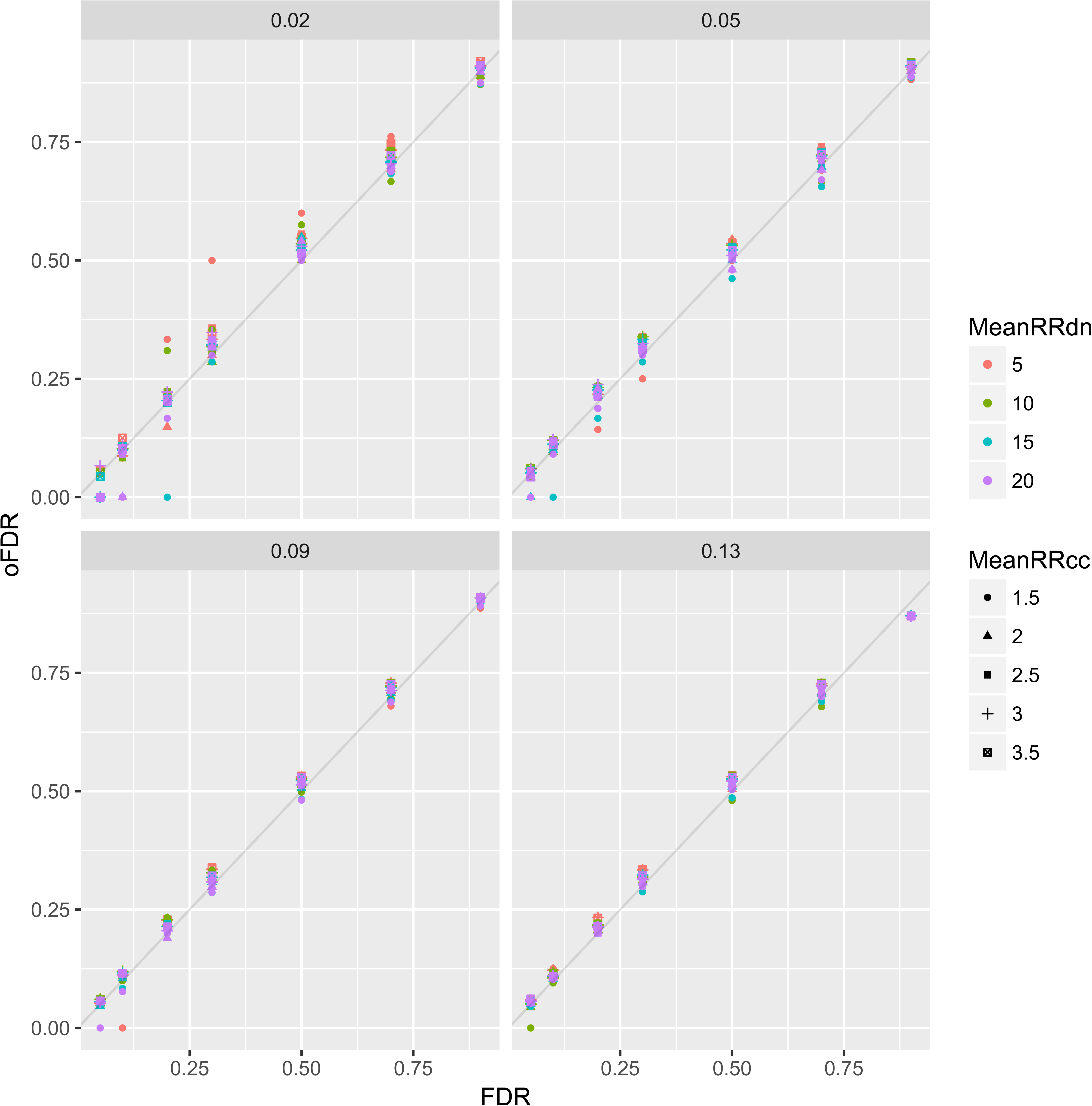
Observed false discovery rates (oFDR) and theoretical FDR (FDR) with different combinations between 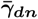 and 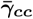. For example, the top left picture shows oFDR and FDR for *π* = 0.02.

### Data for analyses

As shown in Figure S1, the data included samples for schizophrenia (SCZ) and the four NDDs (ID, ASD, DD, and EPI). Variants were annotated as loss-of-function (LoF), missense damaging (MiD), missense (non-damaging unless otherwise stated), silent within frontal cortex-derived DHS peaks (silentFCPk), or silent (not silent-FCPk).

#### SCZ

We applied `extTADA` to the largest available DN and CC SCZ whole exome sequence (WES) data, for inference of rare variant genetic architecture parameters and for genic association. In total, 6,699 cases, 13,028 controls, 1077 trio/quad families were analyzed (Table S1). Primary analyses included three variant categories for DN data: LoF, MiD and silentFCPk, and a single category of CC singletons [5, 7] not present in the Exome Aggregation Consortium (ExAC) [27] (termed NoExAC): LoF+MiD. An array of secondary `extTADA` analyses were conducted to help validate and dissect our results.

*De novo* mutations and case-control variants were tested to select classes and samples for the `extTADA` pipeline. For *de novo* mutations, we calculated the sample-adjusted ratios of mutation counts between 1,077 DN cases and 731 DN controls (Table S1). Similar to [25], the highest ratio was observed for silentFCPk (2.57), followed by MiD (2.3), LoF (1.83), and missense and silent (*∼* 1.3) mutations (Figure S9). Three classes (LoF, MiD and silentFCPk) were used in `extTADA` analyses.

Since currently `extTADA` requires integer counts data, adjustment for ancestry and technical covariates is not possible; therefore, CC data were restricted to more homogeneous population samples (see Supplementary Methods). First, for the 4,929 cases and 6,232 controls from Sweden population sample, we clustered all cases and controls based on PCA and tested each cluster for case-control differences with and without adjustment for covariates. We carried two clusters forward for analysis (Groups 1 and 3 in Figure S8), one with 3,157 cases and 4,672 controls, and one with 1,091 cases and 1,193 controls. We used only the larger UK population sample from the UK10K project data [8], as it showed comparable case-control differences to the homogenous Sweden samples. As in [7], NoExAC singleton CC variants showed significant case-control differences and InExAC variants did not (Figure S8); therefore, we used only NoExAC CC singletons in primary `extTADA` analyses, however we also used all singletons in a secondary analysis for comparison. LoF and MiD variants showed similar enrichment in our case-control data (Figure S8); therefore, we pooled them in order to maximize the case-control information.

#### NDDs

Sample sizes of these diseases are shown in Table S1 and Figure S1. Numbers of trios ranged from 356 for EPI, 1,112 for ID, 4,293 for DD, 5,122 for ASD. As previously reported (see references in Table S1, these data have strong signals for DN mutations contributing to disease (Table S7). Only ASD data included case-control samples (404 cases, 3,654 controls) from the Swedish PAGES study of the Autism Sequencing Consortium [30] (see Supplementary Methods for details).

### Rare variant genetic architectures inferred by extTADA

#### SCZ

`extTADA` generated joint posterior density samples of all genetic parameters for SCZ (Table 1, Figures 2 and S10). All MCMC chains showed convergence (Figure S11). The estimated proportion of risk genes was 8.01% of the 19358 analyzed genes (1551 genes), with 95% CI (4.59%, 12.9%; 890 to 2500 genes). DN LoF variants had the highest estimated mean RR, 12.25 (95% CI 4.78, 22.22). Estimated mean RRs were 1.22 (95% CI 1, 2.16) for silentFCPk and 1.44 (95% CI 1, 3.16) for MiD. For CC MiD+LoF variants, the two Sweden samples had nearly equal mean RR estimates, 2.09 (95% CI 1.04, 3.54) and 2.44 (95% CI 1.04, 5.73), and larger than that of the UK sample, 1.04 (95% CI 1, 1.19).

**Figure 2.**
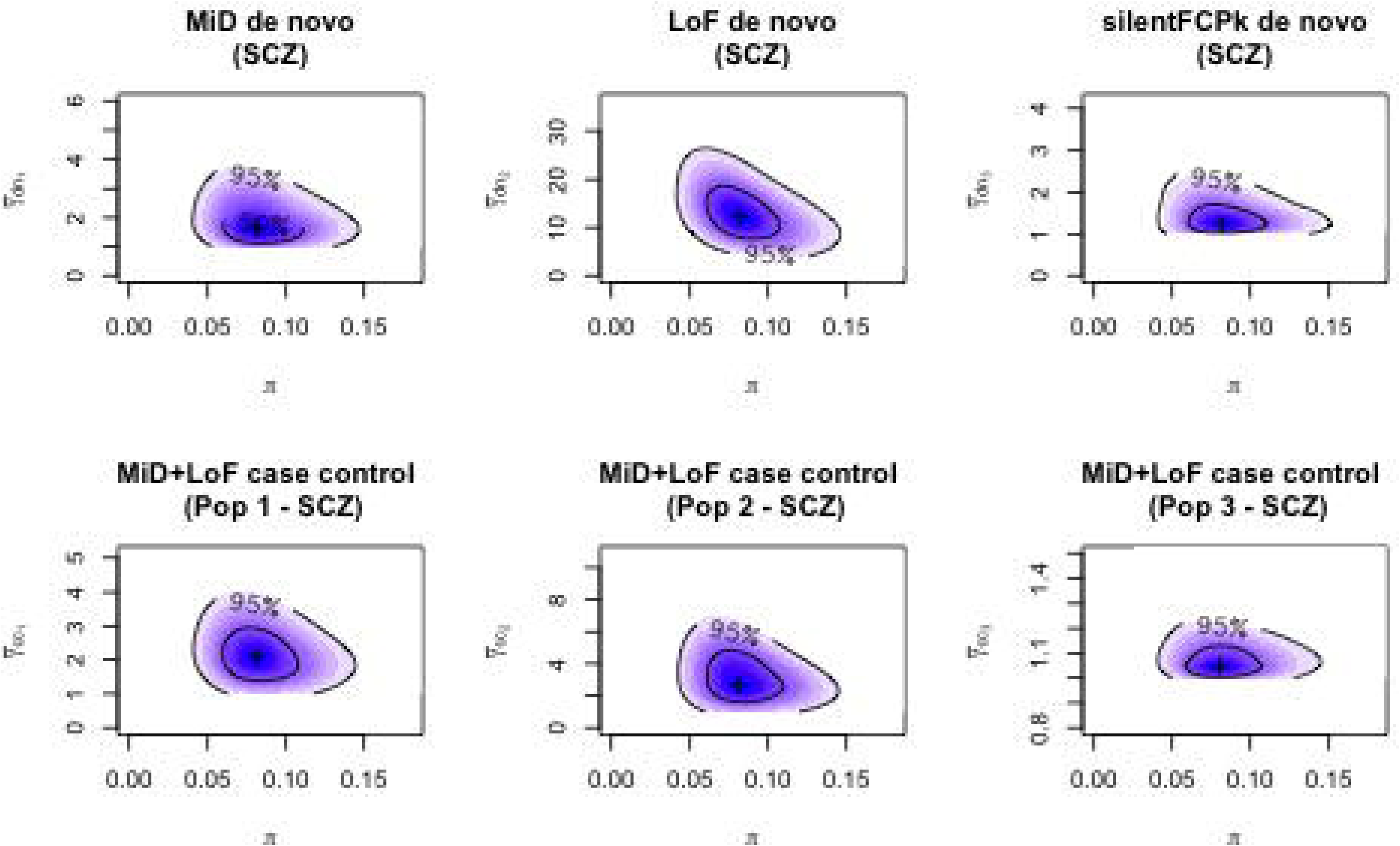
The densities of the proportion of risk genes and mean relative risks for SCZ data. These are obtained after 20,000 iterations of three MCMC chains. The first two case-control populations are derived from the Sweden data set while the third case-control population is the UK population.

**Table 1.** Estimated parameters for *de novo* and case-control SCZ data and four other NDDs: ID, EPI, ASD and DD. These results are obtained by sampling 20,000 times of three MCMC chains. The two last columns show the lower (lCI) and upper (uCI) values of credible intervals (CIs)

| Parameter | Estimated mode | ICI | uCI |
| --- | --- | --- | --- |
| SCZ_pi (%) | 8.01 | 4.59 | 12.9 |
| SCZ_meanRR_silentFCPk_denovo | 1.22 | 1.00 | 2.16 |
| SCZ_meanRR_MiD_denovo | 1.44 | 1.00 | 3.16 |
| SCZ_meanRR_LoF_denovo | 12.25 | 4.79 | 22.22 |
| SCZ_meanRR_MiD+LoF_CCpop1 | 2.09 | 1.04 | 3.54 |
| SCZ_meanRR_MiD+LoF_CCpop2 | 2.44 | 1.05 | 5.73 |
| SCZ_meanRR_MiD+LoF_CCpop3 | 1.04 | 1 | 1.19 |
| ASD_pi (%) | 4.44 | 3.15 | 5.94 |
| ASD_meanRR_MiDdenovo | 3.71 | 2.06 | 8.71 |
| ASD_meanRR_LoFdenovo | 24.56 | 14.27 | 37.44 |
| ASD_meanRR_LoFcc | 4.04 | 2.08 | 8.24 |
| ID_pi (%) | 2.53 | 1.89 | 3.43 |
| ID_meanRR_MiDdenovo | 29.82 | 18.86 | 46.1 |
| ID_meanRR_LoFdenovo | 105.45 | 73.27 | 143.29 |
| DD_pi (%) | 2.84 | 2.29 | 3.45 |
| DD_meanRR_MiDdenovo | 23.42 | 13.97 | 33.97 |
| DD_meanRR_LoFdenovo | 88.32 | 67.54 | 115.09 |
| EPI_pi (%) | 1.14 | 0.52 | 2.1 |
| EPI_meanRR_MiDdenovo | 72.2 | 35.39 | 128.46 |
| EPI_meanRR_LoFdenovo | 89.71 | 45.31 | 169.43 |

To test the performance of the pipeline on individual categories and to assess their contributions to the overall results, we ran `extTADA` separately on each of four single variant classes: silentFCPk, MiD and LoF *de novo* mutations, and MiD+LoF case-control variants (Table S8). All parameter estimates were consistent with the primary analysis, with broader credible intervals. The much larger 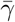 CIs than in integrative analyses demonstrated `extTADA`’s borrowing of information across data types (also observed in simulation, Figure S5).

We also assessed the sensitivity of genetic parameter inference in several secondary analyses. We tested `extTADA` for *de novo* mutations not present in the ExAC database, mutation rates adjusted for the ratio of observed to expected synonymous DN mutations, and alternative model specification of variant annotation categories. Adjusting mutation rates by a factor 0.81, DN mean RR estimates slightly increased as expected, and the estimated proportion of risk genes increased slightly to 9.37% (95% CI 5.47-15.12%), while case-control parameters were highly similar (Table S14). Above we assumed that different case-control population samples may have different mean RRs, which could be due to clinical ascertainment, stratification or population specific genetic architectures. Analysis using a single mean RR parameter for all three case-control samples yielded similar *π* and DNM mean RRs and an intermediate CC MiD+LoF mean RR with relatively narrower credible interval, 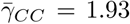 (95% CI 1.08-3.21) (Table S15, Figure S13). Considering all CC singleton variants (not just those absent from ExAC) also generated similar genetic parameter estimates, with slightly lower case-control mean RRs (Table S16).

#### ASD, ID, DD, and EPI

`extTADA` genetic parameter estimates are presented in Table 1 and Figures 3 and S10. MCMC analyses showed good convergence, except for the small sample size EPI data (356 trios compared with over 1000 trios for other diseases). Estimated risk gene proportions (*π*) for the NDDs were lower than that of SCZ. For ASD, estimated *π* was 4.44%, (3.15%, 5.94%), or 859 (610 - 1,150) risk genes, consistent with the result of 550-1000 genes estimated in the original TADA model [16] using only DN LoF data. For DD and ID, *π* estimates were similar, 2.84% or 550 risk genes (2.29%, 3.45%; 443 - 668 genes) and 2.53% or 490 risk genes (1.89%, 3.43%; 366 to 664 genes) respectively, and smaller than that for ASD. The estimated *π* value for EPI, 1.14% or 221 risk genes (0.52%, 2.1%; 101 - 407 genes) was the lowest but with a broad credible interval. The mean RRs of *de novo* mutations in all four NDDs were much higher than those of SCZ, indicating a stronger contribution of *de novo* mutations in these four NDDs. For ASD, estimated mean RRs for de novo mutations were consistent with previous results and much lower than for the other diseases. ID and DD had the highest estimated *de novo* LoF mean RRs, 105.45 (73.27, 143.29) and 88.32 (67.54, 115.09), respectively. Even though the EPI estimated *de novo* LoF mean RR, 89.71 (45.31, 169.43), was similar to those of ID and DD, the estimate for EPI *de novo* MiD mean RR, 72.2 (35.39, 128.46) was somewhat higher than those of other diseases. The previously estimated [67] EPI mean RR of 81 is consistent with the current results, and it will be of interest to see if this result remains consistent in additional data in the future.

**Figure 3.**
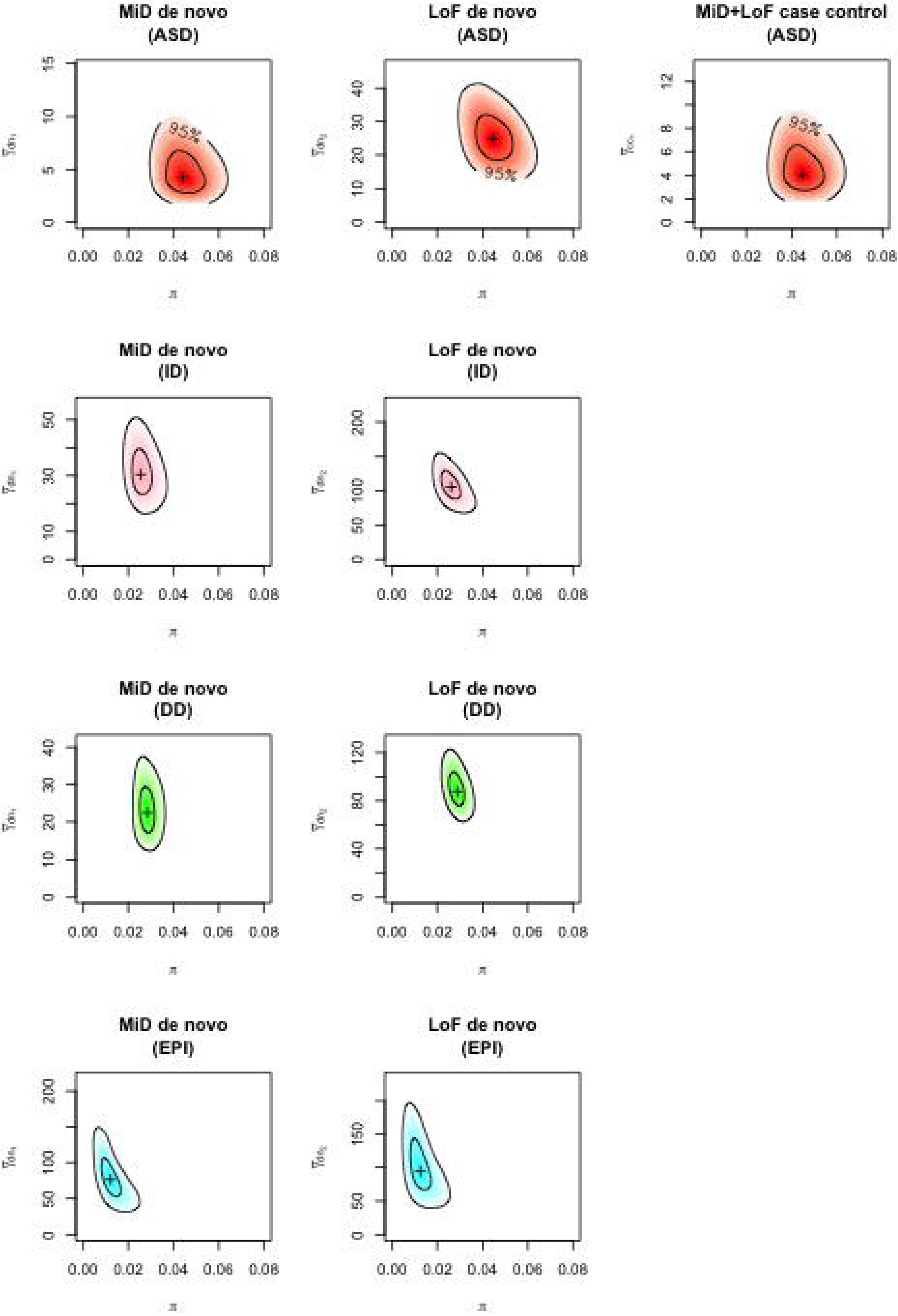
The densities of the proportion of risk genes and *π* mean relative risks (*γ*) for ASD, EPI, ID and DD data. For ASD, there are two *de novo* (dn) classes and one case-control (cc) class. For other diseases, only two *de novo* classes are publicly available for our current study.

### Identification of risk genes using extTADA

#### SCZ

Table S9 includes supporting data as well as association results for SCZ. Four genes achieved PPA *>* 0.8 and FDR *<* 0.1 (*SETD1A, TAF13, PRRC2A, RB1CC1*). Two genes *SETD1A* (FDR = 0.0033) and *TAF13* (FDR = 0.026) were individually significant at FDR *<* 0.05. *SETD1A* has been confirmed as statistically significant in previous studies [8, 25], while *TAF13* was only reported as a potential risk gene in the study of [6]. However, FDR was high (0.74) for the gene *RBM12* which was reported as a risk gene for psychosis by [9]. If we increase the FDR threshold to 0.3 as in a recent ASD study using TADA [30], we identify 24 candidate SCZ risk genes (*SETD1A, TAF13, RB1CC1, PRRC2A, VPS13C, MKI67, RARG, ITSN1, KIAA1109, DARC, URB2, HSPA8, KLHL17, ST3GAL6, SHANK1, EPHA5, LPHN2, NIPBL, KDM5B, TNRC18, ARFGEF1, MIF, HIST1H1E, BLNK*). Of these, *EPHA5, KDM5B* and *ARFGEF1* did not have any *de novo* mutations (Table S9). We note that still more genes show substantial support for the alternative hypothesis over the null the model [68] (58 genes with PPA *>* 0.5, corresponding to BF *>* 11.49, FDR *<* 0.391; Table S9). We note that the secondary analyses slightly impacted support for individual genes (Tables S17, S16, S15).

#### NDDs

The `extTADA` risk gene results of the four disorders ID, DD, ASD and EPI are presented in Tables S10, S11, S12 and S13. With FDR *<* 0.05, there were 56, 160, 49 and 9 significant genes for ID, DD, ASD and EPI, for FDR *<* 0.1, there were 69, 196, 64 and 10 significant genes.

Genetic parameters inferred after adjusting mutation rates for observed silent *de novo* rates are presented in Table S14. For ASD, ID and EPI, the proportions of risk genes were higher than the results of primary analyses because adjustment ratios were less than 1. As a result, the number of significant genes also increased with different FDR thresholds. For DD, the adjustment ratio was *>* 1 (1.16), and the number of significant genes decreased (134 genes with FDR *<* 0.05). 72/134 genes were not among the 93 DD genes reported in the previous study [69], 33 of which were in the list of curated DD genes [70].

#### Novel significant genes in ID and DD

Results of other *de novo* mutation methods using these same data have been recently reported [40, 69]; nevertheless, `extTADA` identified novel genes with strong statistical support from these recent data.

For ID, we found 56 and 69 genes with FDR *≤* 0.05 and 0.1, respectively. We compared these results with the risk gene list of [40], which included previously reported and novel ID genes. 14 of 56 FDR *≤* 0.05 genes (*AGO1, AGO2, ATP8A1, CEP85L, CLTC, FBXO11, KDM2B, LRRC3C, MAST1, MFN1, POU3F3, RPL26, TNPO2, USP7*) were not on the list. Of the 14 genes, six (*AGO2, CEP85L, CLTC, FBXO11, MFN1, TNPO2*) were strongly significant (FDR *<* 0.01); these were genes hit by two or three MiD or LoF *de novo*s but were not identified by the analyses of [40]. We sought to learn more these 14 genes. First, pLI, RVIS information of these genes were obtained. Two genes (*LRRC3C, POU3F3*) had no pLI or RVIS information; therefore, only 12 other genes were tested. The median of pLIs was 1 (observed 1; simulated data: *µ* = 0.11, *σ* = 0.17, *z* = 5.08, empirical p *<* 9.99e-05). In addition, 9 genes (*AGO1, AGO2, ATP8A1, CLTC, FBXO11, KDM2B, MAST1, TNPO2, USP7*) had pLI = 1, and one gene (*RPL26*) had pLI = 0.916. The median of RVISs were −1.49 (observed −1.49; simulated data: *µ* = −0.014, *σ* = 0.21, *z* = −7.03, empirical p *<* 9.99e-05). Second, we looked for these genes in the latest list of curated DD genes released on 18 May, 2017 ([70], accessed on 08 June, 2017). *CLTC* and *FBX011* were in the list. Removing these two genes and recalculating p values as above, pLI information was still highly significant (observed 1, simulated data: *µ* = 0.3, *sd* = 0.39, *z* = 1.7, empirical p was *<* 9.99e-05), and RVIS information was not much different (observed −1.48, simulated data: *µ* = −0.01, *σ* = 0.23, *z* = −6.26, empirical p *<* 9.99e-05).

For DD, there were 160 and 196 genes with FDR *≤* 0.05 and 0.1 respectively. We only focused on the comparison between our results and the 93 significant genomewide results reported by the DD study [69]. Only 52 of 160 FDR *≤* 0.05 genes were among the 93 genes; 98 genes are novel. The 98 genes also included *QRICH1* (FDR = 3.15e-05) which was reported as a suggestive DD gene [69]. Similar to ID, the total MiD+LoF *de novo* counts of these 98 genes were not high (between two and six). Surprisingly, 54 of the 98 novel genes were strongly supported in our results (FDR *<* 0.01). We assessed the known DD genes in the 93 genes with FDR *>* 0.05 and saw two common reasons for the differences. The first was that our missense damaging counts were lower than missense counts of the previous study. It was expected because seven methods were used to define missense damaging mutations. The second reason was that `extTADA` only used the data from 4293 trios while [69] meta-analyzed from multiple studies.

We sought to validate the large number of novel significant DD genes compared with [69] in the same data. First, we compared enrichments of our candidate gene sets in known DD genes and in our novel DD genes. We found that many of the same gene sets were significantly enriched in both previously known and our novel DD genes, with very strong concordance across gene sets (Figure S12). 92/98 novel DD genes had pLI and RVIS information. The median pLIs was 0.997 (observed 0.997; *µ* = 0.033, *σ* = 0.036, *z* = 26.46, empirical p *<* 9.99e-05). The median of RVISs was −0.92 (observed −0.92, simulated data: *µ* = −0.02, *σ* = 0.07, *z* = −11.86, empirical p was *<* 9.99e-05). We also found that 43 of the 98 novel DD genes occur in the latest list of curated DD genes (described above) showing that `extTADA` was able to detect DD genes later identified in other studies. 50 of the 55 novel genes not in the curated DD genes list of had pLI/RVIS information. The median of the 50 pLI values was 0.9415 (observed 0.94, simulated data: *µ* = 0.045, *σ* = 0.064, *z* = 13.95, empirical p was *<* 9.99e-05). The median of RVISs was −0.72 (observed −0.72, simulated data: *µ* = −0.01, *σ* = 0.10, *z* = −6.87, empirical p *<* 9.99e-05). And 23/50 genes were inside the 95 connected genes above. Finally, we used GeNets with the InWeb protein-protein interaction (PPI) network [63] to test the connections between the 98 genes and 93 known genes (191 genes in total). 94 genes (46 known and 48 novel) out of 191 were connected to 8 communities (overall p = 0.006, and community connectivity p *<* 2e-3) (Figure 5).

**Figure 4.**
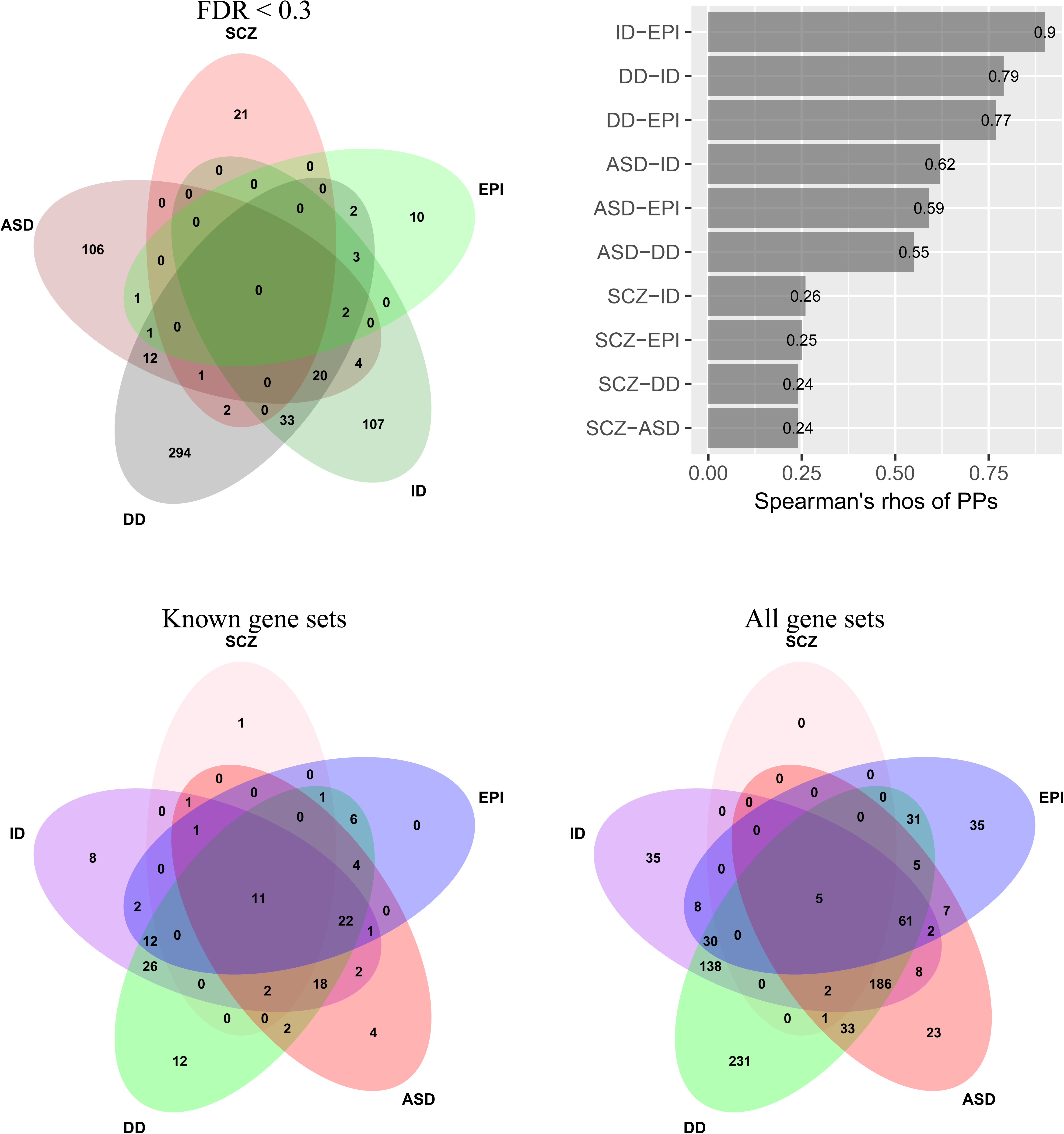
Comparing between five conditions. Top left picture shows overlaps of top significant genes (FDR *<* 0.03). The top right picture describes the correlations of posterior probabilities (PPs) between SCZ, ASD, DD, ID and EPI (all p values *<* 0.0001). These results are calculated by using PPs from `extTADA`. The two bottom pictures show overlaps of significant gene sets in SCZ, ASD, EPI, DD and ID. These results are for 185 and 1879 gene sets respectively.

**Figure 5.**
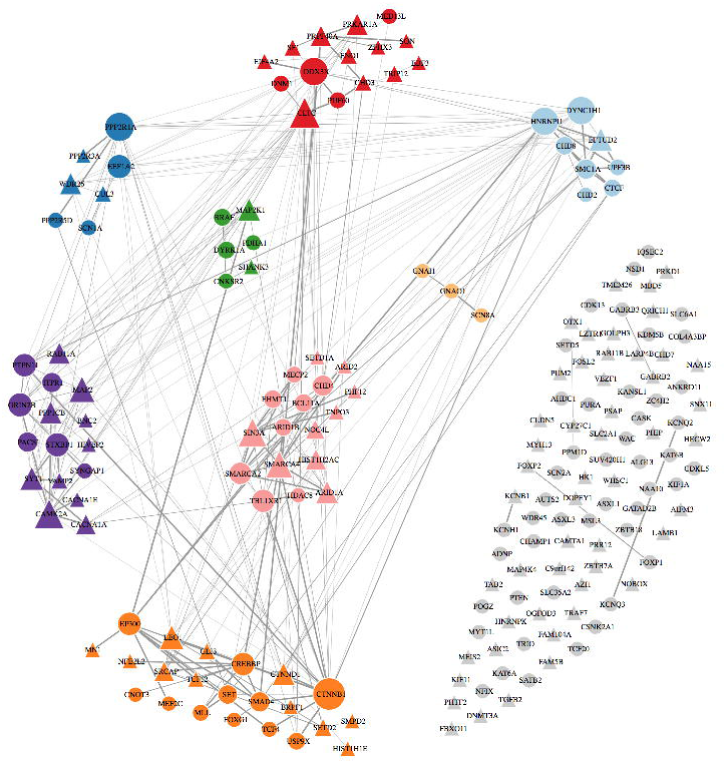
GeNets network analysis for DD significant genes (p *<* 2e-03). These are 93 genome-wide significant genes from [69] and 98 significant genes (FDR *<* 0.05 from `extTADA`) not in the 93 genes. Triangular shapes are the 98 novel genes from `extTADA`.

### Power analysis under inferred genetic architecture

We simulated risk gene discovery using `extTADA` using the genetic architecture of SCZ inferred from the current data (Figures 6 and S14). Samples sizes from 500-20,000 trio families and 1,000-50,000 cases (controls = cases) were simulated as in our validation analyses, using parameters from the posterior distribution samples given the SCZ data, and the number of risk genes with *F DR ≤* 0.05 ranged from 0 to 238. Based on this analysis, we expect *>* 50 risk genes with total sample sizes of trio families plus case-control pairs *γ* 20,000. The results imply that, assuming sequencing costs are proportional to the number of individuals, generating casecontrol data is more efficient than trio data despite the larger relative risks of *de novo* mutations.

**Figure 6.**
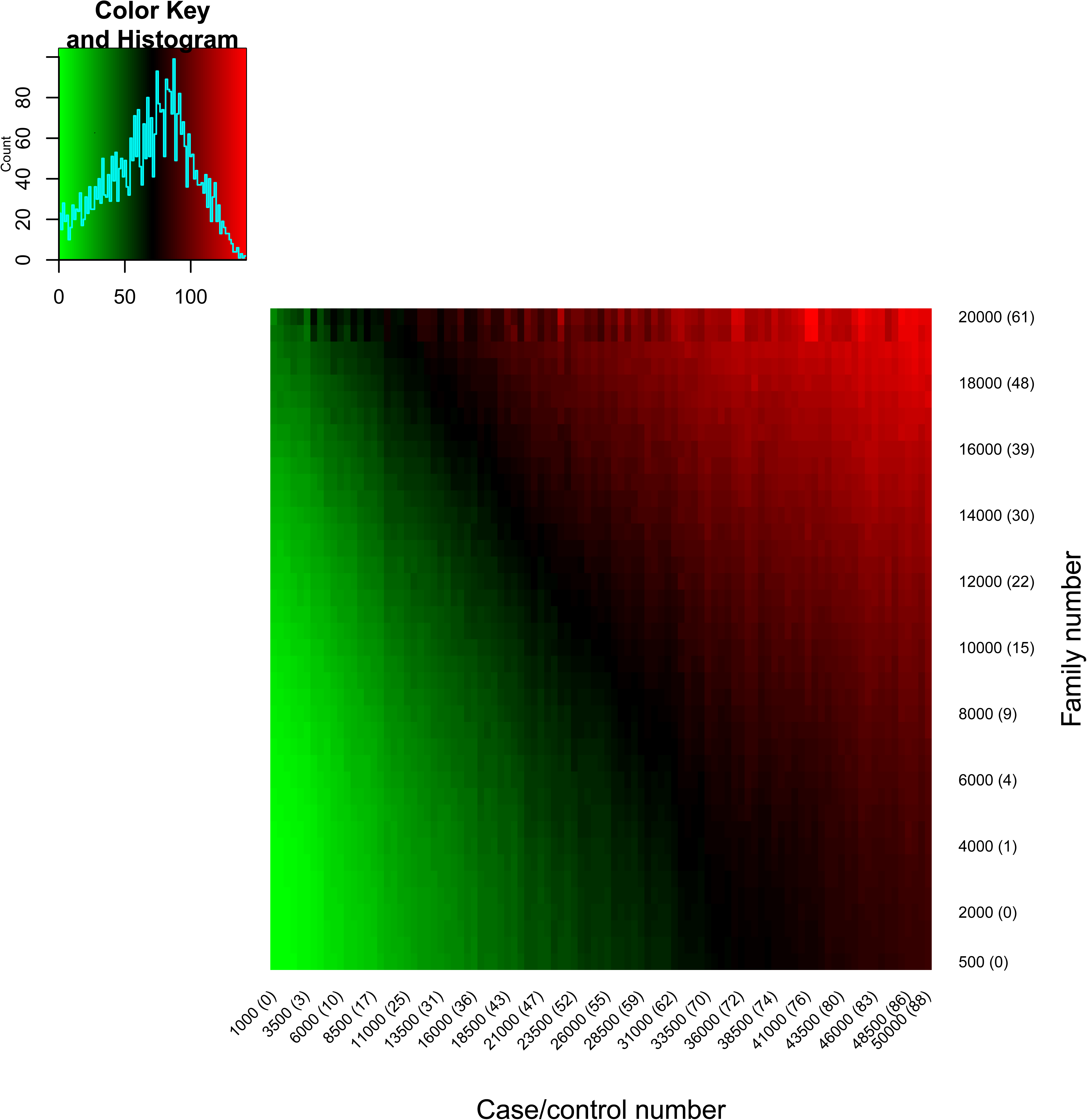
Number of risk genes with different sample sizes based on genetic architecture predicted by extTADA. Case/control number is only for cases (or controls); therefore if Case/control number = 10,000 this means total cases+controls = 20,000. The numbers in brackets show risk-gene numbers if we use only case-control data or only *de novo* mutation data.

### Gene-set enrichments

*Known and novel gene sets are enriched in SCZ risk genes from `extTADA`* We tested 185 gene sets previously implicated in SCZ genetics or with strong evidence relevant to SCZ rare variation [5, 7, 15, 41, 67, 38] (Table S2). FDR-significant results (adjusted p *<* 0.05) were observed for 17 gene sets including those previously reported using these data [5, 6, 7] (Table 2). The most significant gene sets were missense constrained and loss-of-function intolerant (pLI09) genes, targets of RBFOX1/3 and RBFOX2 splicing factors, CHD8 promoter targets, targets of the fragile X mental retardation protein (FMRP) and CELF4 targets (all p *<* 2.0e-04, adjusted p *≤* 7.13e-03, Table 2). Genes harboring *de novo* SNPs and Indels in DD, and post-synaptic density activity-regulated cytoskeleton-associated (ARC), NMDA-receptor (NMDAR) and mGluR5 complexes were alse enriched. Genes exhibiting allelic bias in neuronal RNA-seq data [38] were also enriched in SCZ `extTADA` results (p = 1.9e-03, adjusted p = 2.58e-2). The two brain RNA-seq coexpression modules derived from hippocampus [47] were also significant, M3 and M13. Finally, significant enrichments were also obtained the mouse mutant gene sets with psychiatric-relevant phenotypes including abnormal emotion or affect behavior, abnormal cued conditioning behavior, abnormal sensory capabilities*|*reflexes*|*nociception (FDR *<* 0.05).

**Table 1.**
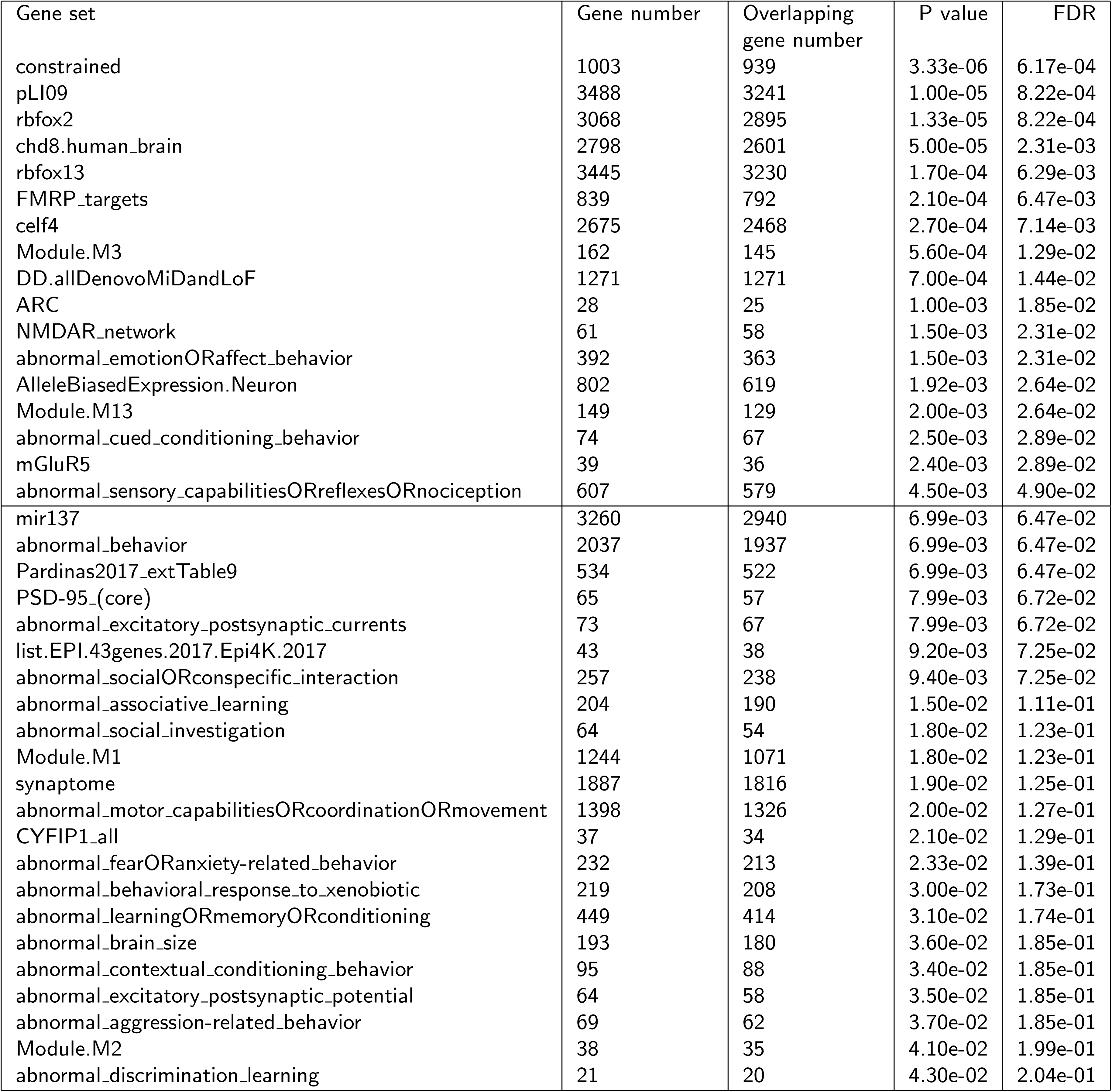
Enrichment of known gene sets from `extTADA` results for SCZ. These P values were obtained by 10,000,000 simulations, and then adjusted by using the method of [57]. The information for these gene sets is summarized in Table S2.

To test more novel gene sets for enrichment in the SCZ `extTADA` results, we tested 1,902 gene sets from several data bases, and FDR-adjusted for the full set of 1,717 + 185 = 1,902 gene sets tested (Tables S18). We used GO, KEGG, REACTOME and C3 sets from MSigDB [71], filtered for sets including greater than 100 genes (see Methods for details). Significant results were observed in 8 gene sets including 6 gene sets in the known gene sets. The top known gene sets still had the lowest p values in these results. We observed significant enrichment of two C3 conserved non-coding motif genesets [72]: GGGAGGRR V$MAZ Q6, genes containing the conserved M24 GGGAGGRR motif; and ACAGGGT,MIR-10A,MIR-10B, including microRNA MIR10A/B targets (Table S18).

#### Multiple gene sets are enriched across NDDs

We saw above that genes containing DN mutations in several of the diseases studied here are enriched in SCZ `extTADA` results. We therefore tested gene set enrichment in the four NDDs and combined this information with the SCZ gene-set information above (Tables S19 and S20). Of the 185 known or strong-candidate gene sets tested in SCZ, 106, 116, 68 and 60 gene sets were significant (FDR *<* 0.05) for ID, DD, ASD and EPI respectively. There were 11 gene sets significant across all five diseases: constrained, PLI09, rbfox2/13, FMRP targets, CELF4, ARC, NMDAR network, abnormal emotion*|*affect behavior, abnormal sensory capabilities*|*reflexes*|*nociception abnormal excitatory postsynaptic currents, hippocampus coexpression module M3 [47]. The significant result of genes in M3 replicated the result of [47]. However, we note that many more gene sets were significant across two or more NDDs, but not SCZ (Figure 4). Our broader set of these 1902 gene sets showed a similar pattern of sharing; only five gene sets which were significant (FDR-adjusted p *≤* 0.05) in all five diseases, while many more gene sets were significant across two or more NDDs (Figure 4).

To validate gene-set results above, we tested gene-set enrichment using the number of gene set genes in the `extTADA` top 500 genes. We saw high correlations between the PP-mean based approach above and this approach (Figure S15).

### Network facilitated interpretation of NDD risk genes

*Overlap among NDD `extTADA` results* There was no gene significant across SCZ and the four NDDs with FDR *<* 0.05 or 0.1. Only *SCN2A* was significant across the four NDDs with these thresholds, but not in SCZ (FDR = 0.35). This gene has been reported as a strong risk gene for multiple NDDs (Reviewed in [2]). Only one additional gene, *STXBP1*, was significant across the four NDDs when the threshold FDR increased to 0.3, but it was not significant for SCZ (FDR = 0.9). At FDR *<* 0.3, several genes were shared among two or three NDDs, whereas only three genes were shared between SCZ and any NDD (Figure 4). We also calculated the correlations between risk gene posterior probabilities for all diseases. Interestingly, high correlations were observed for the four NDDs (*ρ >* 0.5) but not for SCZ and the NDDs (*ρ <* 0.3, Figure 4), both for all genes and significant/suggestive genes in any disease. The pattern of sharing of top `extTADA` results across diseases was consistent when examining gene set enrichments (Figure 4).

Given the high level of sharing among neurodevelopmental disease risk genes, and the large number of novel significant genes we identified, we undertook network analyses to assess and interpret the neurodevelopmental disease risk genes. We chose 288 NDDs genes with different FDR thresholds across the four NDDs to balance the number of significant genes. These thresholds were 0.05 for DD, 0.1 for ASD and ID, and 0.5 for EPI.

First, we used GeNets [63] to test for significant connectedness and structure of NDD genes in the InWeb protein-protein interaction (PPI) network. Including 2nd degree indirect connections, the 288 NDD genes were connected with 89 candidate genes to make a network of 377 genes. These 377 genes were connected in 7 communities (subnetworks, C1-C7), including 149 of the 288 NDD genes (overall connectivity p value and connectivity p values for each community *<* 1.3e-5, Figure 7 and Table S21). Canonical pathway enrichments were observed for five communities, suggesting that they are functional distinct. Significant pathways included betacatenin nuclear signaling, transcriptional regulation of white adipocyte differentiation, WNT signalling pathway, circadian clock (C2); release of several neurotransmitters (C3); splicesome (C4); ribosome, 3’ UTR mediated translational regulation (C5); and neurotransmitter receptor biding and downstream transmission in the postsynaptic cell, calcium signaling, post NMDA receptor activation events (C6) (Table S22). Similar results were obtained restricting the network to direct edges only (connectivity p *<* 0.002, Figure S16), although the resulting 12 communities were less functionally distinct in pathway enrichments.

**Figure 7.**
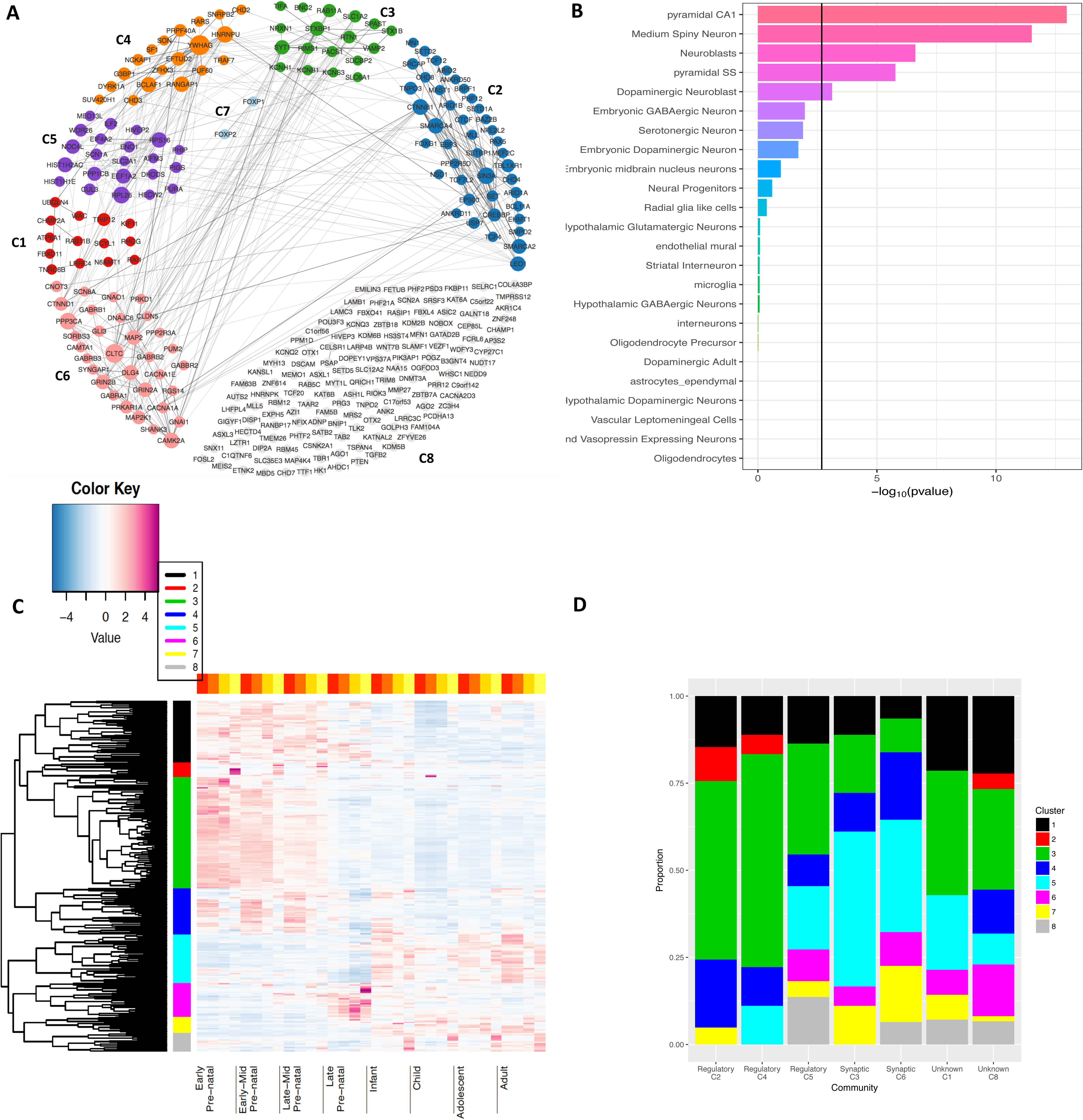
Analyzing results for 288 NDD genes. (A) GeNets results for top 288 NDD genes. 149/288 genes were connected into 7 main communities (colorful genes) and the unconnected genes were put into the 8th community. (B) The enrichment of the 288 genes in different cell types. (C) Grouping the 288 genes to distinct spatiotemporal expression. Genes were clustered into 8 groups using a hierarchical clustering method (color bar). (D) The proportions of different clusters in the 8 communities.

Second, we used mouse single-cell RNA-seq data [53] to test NDD gene enrichments across brain cell types. Significant results were observed for hippocampal CA1 pyramidal cells (p = 1.6e-9), followed by neuroblasts, medium spiny neuron cells, somatosensory pyramidal cells, and dopaminergic neuroblasts (p *<* 6.6e-4, Figure 7). We further tested each GeNets PPI community separately (Figure S17), and found multiple cell types enriched in five communities C2-C6, consistent with their regulatory or synaptic pathway enrichments. Specifically, C2, C4 and C5 were significantly enriched in neuroblasts and neural progenitor cells; C3 and C6 were enriched for pyrimidal CA1 and SS cells (among a few others).

Third, we used BRAINSPAN RNA-seq data to cluster the 288 genes based on their spatiotemporal expression in the developing brain (Figure 7). The genes clustered into 8 groups, and again correlated with PPI communities. Genes in pre-natally expressed groups (Clusters 1,3,4) were over-represented the regulatory communities C2 and C4, p = 3.78e-05. Post-natally expressed groups (Clusters 5,7,8) were in higher proportions in the synaptic communities C3 and C6, p = 1.42e-07.

## Discussion

In this work, we built a pipeline `extTADA` for integrated Bayesian analysis of *de novo* mutations and rare case-control variants in order to infer rare variant genetic architecture parameters and identify risk genes. We applied `extTADA` to available data in schizophrenia and four other neurodevelopmental disorders (Figure S1).

### The extTADA pipeline

`extTADA` is based on our previous work in autism sequencing studies, TADA [16, 30], and conducts fully Bayesian analysis of a simple rare variant genetic architecture model. TADA borrows information across all annotation categories and *de novo* and case-control samples in genetic parameter inference, which is critical for sparse rare variant sequence data. Using Markov Chain Monte Carlo, `extTADA` samples from the joint posterior density of risk gene proportion and mean relative risk parameters, and provides disease-association Bayes factors, posterior probabilities and FDRs for each gene being a risk gene. Inference of rare variant genetic architecture is of great interest in its own right [73], but of course risk gene discovery is a primary objective of statistical genetics. We have shown how the two are not separable, through power analysis of larger sample numbers under the inferred genetic architecture parameters (Figure 6). These analyses, incorporated into `extTADA`, show how study design should be influenced by analysis of currently available data. We hope that `extTADA` (*https://github.com/hoangtn/extTADA*) will be generally useful for rare variant analyses across complex traits.

As in all Bayesian and likelihood analyses, we must specify a statistical model; the true model underlying the data is unknown and could in principle yield different results. This is addressed by analyzing a simple model that can allow illustrative, interpretable results, and by assessing sensitivity to alternative model specifications. `extTADA` uses relatively agnostic hyper-parameter prior distributions (Figure S1), without previously known risk gene seeds. `extTADA` assumes that different variant classes share risk genes such that the mixture model parameter *π* applies to all data types, facilitating borrowing of information across classes. This is supported by convergent *de novo* and case-control rare variant results in SCZ [6, 5, 8, 7] (Table S8); however, some evidence exists for disjoint risk genes for *de novo* vs case-control protein-truncating variants e.g. in congenital heart disease (CHD) [74]. We assume Poisson distributed counts data and Gamma distributed mean relative risks across genes for analytical convenience. The Poisson is likely to approximate genetic counts data well [16], assuming linkage disequilibrium can be ignored and that stratification has been adequately addressed. Poisson *de novo* counts further assume known mutation rates; in our data, mutation rate adjustment for silent *de novo* rates was actually anti-conservative (except for DD). Differences between *de novo* studies are not unlikely even though previous studies of [30, 8] did not adjust mutation rates to account for it. Additional limitations include that we are using public data sets from different sequencing centers, with different technologies and coverages. Thus, although we developed `extTADA` to utilize summary counts data, care must be taken to avoid sample heterogeneity particularly when individual level data are not available. The ability to incorporate covariates, perhaps by modeling Gaussian sample frequency data, would be an important further extension of TADA-like models.

### Insights for schizophrenia

The current study generally replicated previous studies and generated new insights for schizophrenia. In this study, we described in detail the rare variant genetic architecture of SCZ. It appears more complex than those of ASD, ID, DD and EPI; the estimated number of SCZ risk genes, *∼* 1, 551, is higher than those of the four other NDDs, and their relative risks are weaker (Figures 2 and 3, Table 1). Based on our inference, we showed that tens of thousands of samples are required to identify many rare variant risk genes (*≥* 50) [73], and that, in contrast to autism studies [30, 16], case-control studies may be more efficient than trio studies in risk gene identification. We found that *SETD1A* [8, 25] is the most significant gene across analyses (FDR *∼* 1.5 *×* 10^-^3^^), and that *TAF13* [6] is FDR significant. Of two genes with 0.05 *<* FDR *<* 0.1, rare duplications covering *RB1CC1* have been reported in SCZ [75] and in ID and/or DD [76]. Two novel conserved non-coding motif gene sets showing brain specific expression [72] were enriched (Table S18), including targets of the transcription factor MAZ and of microRNAs MIR10A/B. In addition, we see slight overlap between rare and common variant genes [15] (p = 0.007, FDR = 0.06).

### Insights into neurodevelopmental disorders

We used `extTADA` to infer genetic parameters for four other neurodevelopmental disorders ASD, EPI, DD and ID (Table 1, Figure 3). The ASD results of `extTADA` are comparable to previous results [16, 30]. We found lower risk gene proportions particularly for DD and ID, and exceptionally high *de novo* missense damaging mean RR estimated for EPI (also consistent with previous analyses [77]). These parameters facilitate the identification of novel risk genes, particularly for DD. We did not restrict out primary analyses to private DN mutations (not in ExAC) as recently discussed [78]); however, we note that mutation rate calibration might be required for analyses focusing on private mutations. Nonetheless, multiple ID/DD genes discovered in this study are in lists of curated ID/DD genes. In addition, our novel significant genes have similarly high conservation (e.g. pLI, RVIS) as ID/DD genes discovered recently by [40]. This shows that using both private and non-private *de novo* mutations also has the power to obtain significant genes. We also highlight the sharing of risk genes across the neurodevelopmental disorders (Figure 4). Multi-phenotype analyses leveraging this sharing could have higher power to to detect novel risk genes.

We conducted network analyses of 288 top NDD risk genes from `extTADA`. We identified highly significant protein-protein interaction connectivity, and communities differentially enriched for functionally distinct canonical pathways (Figure 7 and Table S22). A substantial number of genes found are synaptic, particularly present in communities C3 (presynaptic) and C6 (postsynaptic). The presynaptic PPI community identified in this study (C3, Figure 7), accumulates genes for which synaptic phenotypes are particularly strong in null mutant mice (*STXBP1, STX1B, SYT1, RIMS1, VAMP2*). *STXBP1*, the only significant gene across the four NDDs (FDR *<* 0.3), is a presynaptic gene involved in preparing synaptic vesicles for regulated secretion (reviewed in [79]). The stxbp1 (munc18-1) null mutant shows a loss of all aspects of synaptic transmission, and spontaneous events [80], the strongest phenotype among all mutants described to date for presynaptic genes. Loss of one copy of the gene in mice leads to subtle synaptic defects [81], a defect more severe in inhibitory neurons than in excitatory neurons [81], therefore implicating excitation/inhibition imbalance, a central aspect in epilepsy pathogenesis, implicated also in autism and schizophrenia [82]. Known clinical features of de novo heterozygous *STXBP* mutations (reviewed in [83]) include severe intellectual disability, seizures and autistic traits [83].

Of the postsynaptic density proteins, C6 includes the prerequisite glutamate-gated ion channel-forming subunit GRIN1 of the NMDA receptor complex. In contrast to AMPA-type glutamate receptor subunits, which are not present, NMDARs are important for Ca-dependent signaling and plasticity processes. The Ca-dependent calmodulin kinase II (CAMK2A) and phosphatase PPP3CA are also identified as NDD risk genes in C6. Other important protein phosphatases are found in different communities, PPP1CB (C5), PPP2R5D (C2). Mutations in these Ca-mediated signaling proteins are well known to affect synaptic plasticity and lead to major neuronal dysfunction [84, 85, 86, 87, 88].

The postsynaptic community C6 also contains the three GABA-binding beta subunits (GABRB1-3) of the GABAA receptor (out of the myriad of GABAA receptor subunit diversity), G-protein coupled receptor signaling (GABBR2, RGS14, GNAO1), cell adherence-mediated signaling (CNNTD1, and CNNTB1 in C2), and the major postsynaptic density protein-interaction scaffold organizing proteins DLG4, SHANK3 and SYNGAP1, mutants of which have been shown to have major impact on synaptic function [89, 90]. Also notable among the 288 NDD risk genes are ion channels with roles in excitability including calcium channel subunits CACNA1A/1E (C6) and the auxiliary calcium channel subunit CACNA2D3 (C8), three pore-forming sodium channel subunits SCN8A (C6), SCN1A (C5), and the well-known strong NDD risk gene SCN2A (C8), and potassium channel subunits KCNQ2/3 (C8) [91]. Finally, transcriptional activator AUTS2 occurs in unconnected C8 and is a candidate for NDDs including ASD, ID and DD [92].

In single-cell RNA-seq data, the top enriched cell types were CA1 pyramidal cells and striatal medium spiny cells, similar to SCZ [53]; but in contrast to SCZ, neuroblasts and neural progenitor cells were also clearly enriched for NDDs. The PPI analysis reveals that different cellular machineries are involved in NDD etiology; in neuroblasts this was driven by PPI communities (C2, C4, C5) enriched in regulatory pathways, while in pyramidal neurons this was driven by the synaptic communities (C3, C6). Finally we note that NDD risk genes clustered by expression across development revealed significant but imperfect correlation with PPI communities, and, together with the occurrence of at least some known interactors across PPI communities (see above), argues that even synaptic proteins confer risk at different stages of development, perhaps through as yet unknown mechanisms.

## Conclusions

We have developed the `extTADA` pipeline and analyzed rare variants in schizophrenia (SCZ) and four neurodevelopmental disorders (NDDs). For SCZ, we generated new insights particularly for rare variant genetic architecture. 98 new significant developmental delay (DD) genes were identified and *in silico* validated. These genes are highly connected with previous DD genes by analysing PPI network, and have similar conservation and gene set enrichments to known DD genes. To better understand NDD genes, we further analyse 288 top NDD genes from `extTADA`. PPI network analysis shows that these genes are strongly connected in functionally distinct subnetworks based on canonical pathway enrichments, single-cell RNA-seq cell types, and developmental transcriptomic data, revealing some of the most important players and processes dysregulated in neurodevelopmental disorders.

## Author’s contributions

Designed the pipeline used in analysis: HTN, EAS. Conceived and designed the experiments: HTN, XH, PFS, EAS. Performed the experiments: HTN, JB, AK, ABMM, EAS. Analyzed the data: HTN, JB, AK. Contributed reagents/materials/analysis tools: HTN, JB, AK, AD, LMH, ABMM, DMR, GG, MF, XX, DP, SL, MV, ABS, JHL, JB, CH, PS, SMP, KL, XH, PFS, EAS. Wrote the paper: HTN, AD, LH, JB, MV, ABS, JHL, XH, PFS, EAS.

## Acknowledgements

This work was supported in part through the computational resources and staff expertise provided by Scientific Computing at the Icahn School of Medicine at Mount Sinai, by NIH grant R01MH105554 to E.A.S, and by NIH grant R01MH110555 to D.P. JB was supported by a grant from the Swiss National Science Foundation. The Sweden exome sequencing data generation and analysis are supported by the Stanley Center for Psychiatric Research and NIH grant R01 MH077139 to C.H, P.S and P.F.S. We are deeply grateful for the participation of all subjects contributing to this research.

## Additional Files

Additional file 1 — Supplementary Information

This file (**AdditionalFile1.pdf**) describes supplementary results, methods, data, figures and short Tables.

*Supplementary Results Supplementary Methods*

*Supplementary Figures*

## Supplementary Figures

This file includes Sup Figures below.

**Figure S1** Workflow of data analysis.

**Figure S2** Comparison between TADA and `extTADA`. They both use the same model for *de novo* data (*x*_*dn*_ and case/control (*x*_*ca*_, *x*_*cn*_) data. `extTADA` combines all categories to obtain parameters and their credible intervals while TADA is based on LoF mutations. `extTADA` uses an approximate model for case-control data, and constrains *β* and 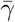 in the estimation process. `extTADA` is designed to work for multiple populations. TADA can be used inside `extTADA`.

**Figure S3** A grid of *β* and 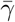 values. Points on the red line are corresponding with the proportion of protective variants less than 2%.

**Figure S4** Correlations between estimated and simulated values for one CC class with different sample sizes. X and Y axes describe simulated (S) and estimated (E) values respectively. The top picture is for mean relative risks (MeanRRs) while the bottom picture is for the proportion of risk genes (*π*). Legends show sample sizes and correlations. These estimated values were averaged across simulation results. Detailed values are presented in Figure S5.

**Figure S5** Correlation between simulated and estimated values for one-category case/control data.

**Figure S6** Correlations between estimated and simulated values for two CC class with different sample sizes. X and Y axes describe simulated (S) and estimated (E) values respectively. A range of mean relative risks for two classes (MeanGamma1 and MeanGamma2) and risk-gene proportions (*π*) were used in the simulation process. Legends show sample sizes and correlations. These estimated values were averaged across simulation results. Detailed values are presented in Figure S7.

**Figure S7** Correlation between simulated and estimated values for two-category case/control data.

**Figure S8** Odds ratios for the analysis of all case-control samples. Top left picture shows odds ratios for all Sweden samples while the three other pictures show odds ratios for three groups after the clustering process. Only group 1 and 3 are used in the current analysis because there are strong differences between results using covariates and not using covariates in group 2. P values were calculated for variants in (InExAC), not in (NoExAC) the ExAC database, and all variants (Both).

**Figure S9** Ratios of *de novo* mutations between SCZ probands and controls (unaffected siblings). “silentFCPk” describes for silent mutations within frontal cortex-derived DHS (silentCerebrumfrontalocPk.narrowPeak). MiD mutations are missense mutations derived from 7 methods.

**Figure S10** Estimated gene counts for all disorders, with 95% CIs. Point sizes are proportional to sample sizes.

**Figure S11** MCMC results for SCZ data. novel genes from `extTADA` results for DD

**Figure S12** The correlation of gene-set p values (values are-log10(p values)) between known and

**Figure S13** SCZ genetic parameters when mean RRs of case-control data are equal.

**Figure S14 Number of risk genes with different sample sizes based on genetic architecture predicted by** `extTADA`. Case/control number is only for cases (with equal controls) (x-axis); each panel shows results for a given number of trios.

**Figure S15** The correlation of gene-set p values (-log(p value)) between mean posterior profitability (meanPP) based method and permutation based methods.

**Figure S16** GeNets InWeb PPI network for 288 NDD genes, with direct edges only.

**Figure S17** Evaluation of enrichment of each community from GeNets results in brain scRNAseq datasets from mouse.

### Supplementary Tables

This part includes Sup Tables below.

**Table S1** *de novo* and case/control data. For ASD studies, [93] integrated previous results in their study; therefore only *de novo* meta data in their study are shown in the table. In addition, for ASD case-control data, only one homogeneous Sweden population from [30] was used. For case-control data of SCZ, after correcting for the population stratification, only 4,248 cases (3,157 + 1,091) + 5,865 (4,672 + 1,193) controls from [7] and 1,353 cases + 4,769 controls from [8] are used in this study.

**Table S2** Abbreviations of known gene sets used in this study.

**Table S3** Parameter information used in all analyses. *N*_*dn*_*, N*_1_*, N*_0_ are sample sizes of families, cases and controls respectively. 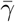 is mean RRs and *β* controls the dispersion of *γ*. 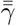 and 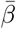 are priors for 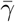 and are set in advance (they are inferred from simulation data). *β* is inferred from the equation 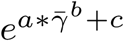 inside the estimation process with a = 6.83, b = −1.29 and c = −0.58.

**Additional file 2 – Long Supplementary Tables Excel file**

**Table S4** Simulated and estimated values of *de novo* (DN) and case-control (CC) parameters. Q50, Q5 and Q95 are for quantile values of 0.5, 0.05 and 0.95 respectively.

**Table S5** Estimated values for the cases in Table S4, for each unique set of parameter values. The first three columns are simulated values. The following columns show estimated *π*, *de novo* mean relative risk (dn RR) and case-contorl (cc) RR; for each parameter, shown are median (Q50) and 5th and 95th %-iles (Q5 and Q95) estimates over 100 simulation replicates.

**Table S6** Estimated values in the case *π* = 0 and 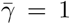. The first two columns are 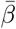 values (prior information of 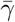: 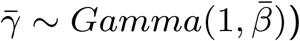). The third to the seventh columns are genetic parameters estimated from `extTADA`. Next columns are the number of risk genes estimated with the corresponding FDR values in the header.

**Table S7** *de novo* mutation counts of categories and their mutation counts per sample size for schizophrenia (SCZ), autism spectrum disorder (ASD), epilepsy (EPI), intellectual disorder (ID) and developmental disorder (DD).

**Table S8** Genetic parameters for SCZ data if single class is used in the analysis.

**Table S9** `extTADA` results of SCZ risk gene identification. (See Long Supplementary Tables xlsx download.)

**Table S10** `extTADA` risk gene identification results of ID data. (See Long Supplementary Tables xlsx download.)

**Table S11** `extTADA` risk gene identification results of DD data. (See Long Supplementary Tables xlsx download.)

**Table S12** `extTADA` risk gene identification results of ASD data. (See Long Supplementary Tables xlsx download.)

**Table S13** `extTADA` risk gene identification results of EPI data. (See Long Supplementary Tables xlsx download.)

**Table S14** SCZ and NDD genetic parameters after adjusting mutation rates.

**Table S15** Estimated genetic parameters for SCZ data with the same mean RRs for case-control data.

**Table S16** SCZ genetic parameters using all variants in and not in ExAC database (InExAC + NoExAC).

**Table S17** `extTADA` results of SCZ risk gene identification after adjusting mutation rates. (See Long Supplementary Tables xlsx download.)

**Table S18** Enrichment of gene sets from different databases with SCZ genes from `extTADA` results. These p values were obtained by 10,000,000 simulations, and then adjusted by using the method of [57]. (See Long Supplementary Tables xlsx download.)

**Table S19** The p values of enrichment tests for known gene sets in SCZ, DD, ID, ASD and EPI. (See Long Supplementary Tables xlsx download.)

**Table S20** The p values of enrichment tests for all gene sets in SCZ, DD, ID, ASD and EPI. (See Long Supplementary Tables xlsx download.)

**Table S21** Community memberships of the GeNets InWeb PPI 288 NDD genes network.

**Table S22** Enrichment results of GeNets. These are enrichment results of 6 communities obtained from GeNets.

